# Fish MicroRNA Responses to Thermal Stress: Insights and Implications for Aquaculture and Conservation Amid Global Warming

**DOI:** 10.1101/2024.11.22.624945

**Authors:** Ting Lin, Madhava Meegaskumbura

## Abstract

Under the backdrop of global warming, heat tolerance emerges as an important physiological trait governing the distribution and survival of fish species worldwide. While knowledge on fish heat tolerance and stress has progressed from behavioral studies to transcriptomic analyses, our understanding at the transcriptomic level remains limited. Recently, the highly conserved nature of microRNAs (miRNAs) has introduced new ways of explaining molecular mechanisms of heat stress in fish. Here, we systematically review current research across three main reference databases to explain the universal responses and mechanisms of fish miRNAs to heat stress. An initial screen of 569 articles yielded 13 target papers for comprehensive analysis. Among these, at least 214 differentially expressed genes (DEGs) were identified, with 15 DEGs recurring in at least two studies (12 upregulated, 13 downregulated). The 15 recurrent DEGs were subjected to DIANA mirPath v.3 utilizing the microT-CDS v5.0 database to identify potential target genes. The results indicate that multiple miRNAs target genes, forming a complex network to regulate glucose and energy balance metabolism, maintain homeostasis, and modulate inflammation and immune disorders. Notably, *miR-1*, *miR-122*, *let-7a*, and *miR-30b* are identified as potential biomarkers of heat stress in fish. Due to their high conservation across species, these miRNAs could be used to monitor the health of wild fish populations, improve selective breeding programs in aquaculture, and guide conservation strategies for species that are vulnerable to climate change. This review provides a framework for further investigation into the molecular mechanisms of fish heat tolerance.

## Introduction

Temperature is a critical abiotic factor that regulates fish physiology and ontogeny, profoundly influencing survival, feeding patterns, reproductive cycles, and locomotion (Handeland, Imsland, & Stefansson, 2008; Naz, Iqbal, Khan, Chatha, & Naz, 2023; Watson et al., 2018). Subtle fluctuations in aquatic temperatures can affect fundamental physiological processes such as respiration, metabolism, and digestion (Y. Chen et al., 2023). Although fish exhibit physiological plasticity, enabling them to acclimate within certain thermal limits (Soyano & Mushirobira, 2018), drastic temperature changes can act as stressors, compromising fish health when these variations approach or exceed species-specific thermal tolerance thresholds (Takegaki & Takeshita, 2020). Stress, a state triggered by external factors and mediated through the endocrine system (Gorissen & Flik, 2016), can induce adaptive physiological or behavioral changes. However, prolonged or repeated stress leads to maladaptive outcomes, impairing growth, reproduction, and immune function, ultimately threatening survival (Barton, 2002).

In the context of escalating climate change, where seasonal water temperatures are rising and extreme thermal fluctuations are becoming more frequent (Vasseur et al., 2014), global warming exacerbates the occurrence of marine heatwaves (Froehlich, Gentry, & Halpern, 2018; Vasseur et al., 2014). Notably, rather than gradual increases in average temperatures, extreme events may be more critical in determining population dynamics (Sandblom et al., 2016). According to the IPCC, global temperatures are expected to rise by 1-4°C by the end of the century (Barbarossa et al., 2021), placing 20-40% of fish species at risk depending on their specific thermal tolerances (Dahlke, Wohlrab, Butzin, & Pörtner, 2020). The variation in tolerance levels is shaped by species’ evolutionary histories, which define their ecological niches, distribution ranges, and resilience to heat stress (Ackerly, 2003; Brauer et al., 2023; Dahlke et al., 2020). For instance, largemouth bass (*Micropterus salmoides*) thrive at 26-29°C (Díaz et al., 2007), whereas turbot (*Scophthalmus maximus*) prefer 13-17°C with a lethal limit of 26-28°C (Jia et al., 2020). Thermal tolerance can vary across life stages even within the same species, with spawning adults and embryos often more vulnerable than larvae and non-reproductive adults (Dahlke et al., 2020). Fish inhabiting tropical, polar, and cave ecosystems or other unique habitats tend to have narrower thermal tolerances, making them particularly susceptible to heat (Kevin T. Bilyk & Devries, 2011; K. T. Bilyk & Sformo, 2021; Peck, Morley, Richard, & Clark, 2014; Tabin et al., 2018). The threat of heat stress is thus unprecedented for global fisheries and poses significant risks to fish biodiversity.

Fish responses to stress occur in three stages: hormonal surges (cortisol, catecholamines) (Shahjahan et al., 2022), metabolic shifts (increased glucose and lactic acid) (Farag et al., 2021; Shahjahan et al., 2022), and physiological or behavioral changes (e.g., growth inhibition, altered behavior, increased disease susceptibility) (Schreck & Tort, 2016). Recent research has taken a molecular approach to understanding stress responses, with transcriptomic analyses providing insights into how fish respond to environmental changes (Raza et al., 2022). MicroRNAs (miRNAs), small non-coding RNAs of ∼22 nucleotides (Tanase, Ogrezeanu, & Badiu, 2012), play a critical role in regulating cellular stress responses by modulating gene expression (Babar, Slack, & Weidhaas, 2008). These miRNAs form RNA-induced silencing complexes (RISC) with AGO proteins, influencing the degradation or translation of target mRNAs, depending on sequence complementarity (Bartel, 2004). miRNAs are involved in critical cellular processes such as proliferation, differentiation, apoptosis, and immune responses, helping maintain homeostasis (Bartel, 2004; C. Z. Chen, Li, Lodish, & Bartel, 2004; Leung & Sharp, 2010; Pedersen et al., 2007). Their differential expression in response to environmental stress is crucial for adaptation (Noman et al., 2017).

Microarray and high-throughput sequencing technologies have advanced our understanding of the physiological pathways mediating environmental adaptation, which offer insights into how organisms tolerate abiotic stress (Herkenhoff et al., 2018). In the context of heat stress, heat shock proteins (HSPs), particularly Hsp70, have emerged as critical players in enhancing thermal resilience across various species, including fish (Kroemer, 2001; F. Wang et al., 2015). For instance, *ssa-miR-301a-3p* has been shown to enhance heat tolerance in rainbow trout by targeting hsp90b2 (Z. Liu, Ma, Kang, & Liu, 2022), while *miR-122* and *miR-30b* in carp are downregulated in response to temperature shifts, regulating lactic acid metabolism through their targets LDH-b and GRHPR (Sun, Liu, et al., 2019). In turbot, heat-responsive genes like ZNF469 and MAGI2 have been proposed as biomarkers for thermal stress (Ising & Holsboer, 2006). However, despite emerging knowledge, our understanding of fish miRNA responses to thermal stress remains incomplete, with many aspects of their role in heat stress regulation yet to be elucidated (Herkenhoff et al., 2018).

Given the increasing frequency of extreme weather events due to climate change, understanding fish thermal adaptation is crucial for conservation efforts and aquaculture development. While research has advanced from behavioral and physiological studies to genetic and molecular approaches, fish species exhibit varied responses to thermal stress, complicating generalizations (Barrett et al., 2011).. For example, rainbowfish populations with higher genetic diversity demonstrate greater adaptability to temperature fluctuations (Brauer et al., 2023). Despite these differences, miRNAs, which are highly conserved across species, offer valuable insights into the universal mechanisms underlying thermal stress responses (Ha & Kim, 2014). Here, we review miRNA expression in teleost fish following heat tolerance experiments to identify biomarkers for thermal stress to facilitate the development of miRNA-based biomarkers that could monitor the health of wild fish populations, enhance selective breeding programs in aquaculture, or inform conservation strategies for vulnerable species.

## Materials and methods

### Review Paper Selection Criteria

To assess miRNA responses to thermal stress in fish, we conducted a systematic review using predefined keywords across three academic databases: (PubMed, Web of Science, Scopus). The following keywords, “fish,” “thermal stress,” “heat shock,” “miRNA,” and “small RNA sequencing,” were used to identify relevant literature (Supplementary Table 1). The search was limited to peer-reviewed articles published between 1993-2024.

From the initial search, 681 results were retrieved. After removing 43 duplicate papers using Excel’s find-and-replace function, 638 unique records remained. Titles and abstracts were then screened based on predefined inclusion criteria: studies had to (1) focus on thermal stress in fish, (2) use miRNA as a primary molecular marker, and (3) involve experimental research. Studies were excluded if they did not involve fish or were review articles, conference proceedings, or non-peer-reviewed literature. Following this screening, 13 research papers were deemed suitable for full-text analysis and inclusion in our review (Fig. 1) (Supplementary Table 2).

**Fig. 1.**
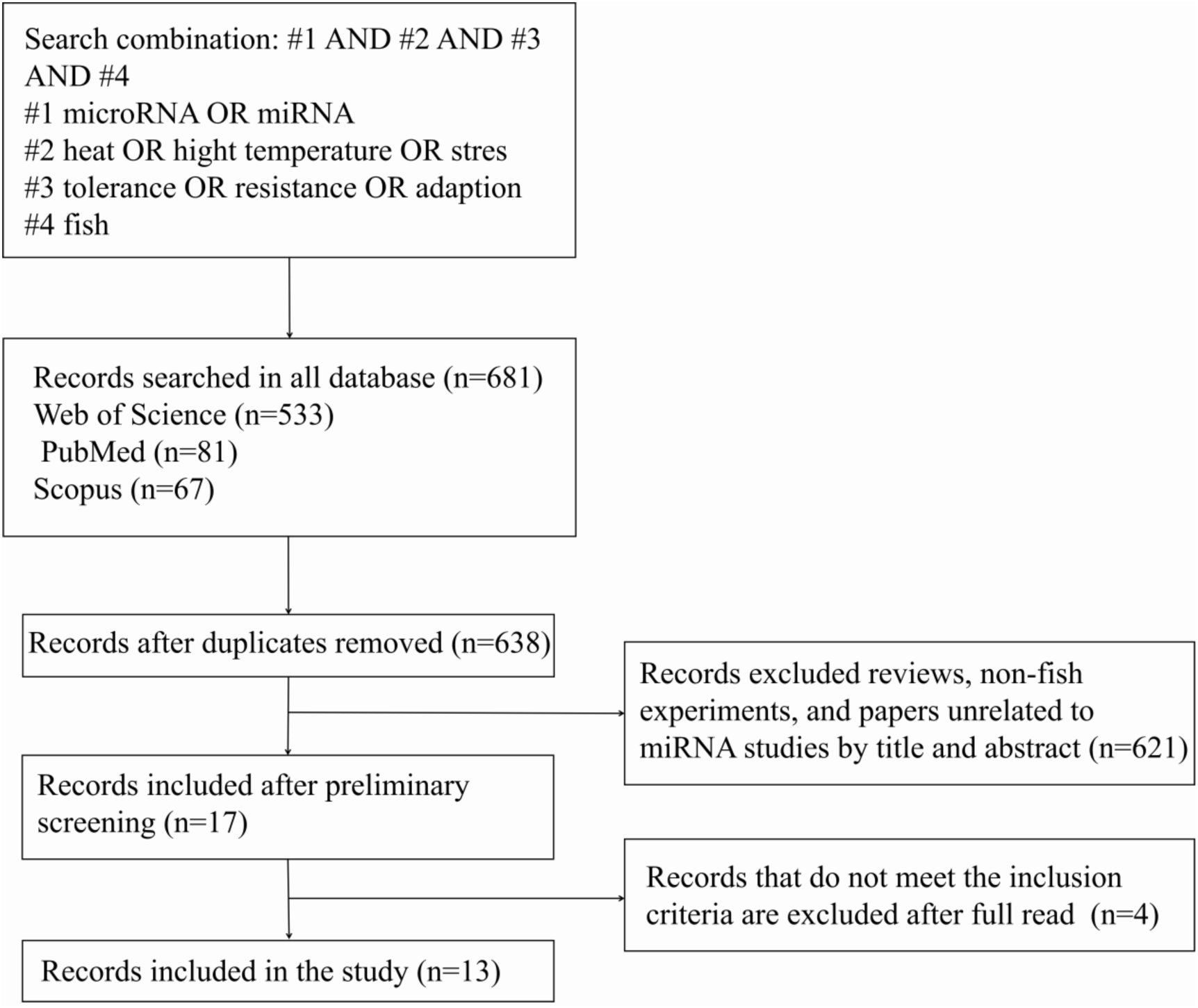
Systematic literature search and screening process for identifying studies on the role of microRNAs in fish heat stress tolerance and adaptation. The flowchart illustrates the systematic process used to identify relevant studies. The search was conducted across three databases—Web of Science (n = 533), PubMed (n = 81), and Scopus (n = 67)—using a search string that combined microRNA or miRNA (#1) with terms related to heat stress or high temperature (#2), tolerance or adaptation (#3), and fish (#4). A total of 638 records were retrieved, with duplicates removed. After preliminary screening based on title and abstract, 17 records were selected, excluding reviews and non-fish or irrelevant miRNA experiments (n = 621). Following a full-text review, 13 studies met the inclusion criteria for the final analysis, and four were excluded.

The selected studies investigated miRNA expression in seven fish species: *Cyprinus carpio* (normal carp) (Sun, Liu, et al., 2019; Sun, Zhao, et al., 2019), *Trematomus bernacchii* (Emerald rockcod) (Vasadia, Zippay, & Place, 2019), *Oncorhynchus mykiss* (rainbow trout) (J. Huang et al., 2018; T. Huang et al., 2022; Z. Liu et al., 2022; Ma et al., 2019; Zhao et al., 2023), *Gymnocypris przewalskii* (Tibetan naked carp) (C. Zhang, Tong, Tian, & Zhao, 2017), *Oreochromis niloticus* (tilapia) (Bao et al., 2018; Qiang, Bao, et al., 2017), *Alosa sapidissima* (American shad) (Y. Liu et al., 2024), and *Gadus morhua* (Atlantic cod) (Bizuayehu, Johansen, Puvanendran, Toften, & Babiak, 2015). The experimental subjects included adult fish, juveniles, and larvae, and organs studied encompassed the liver, gills, head kidney, brain, eye, muscle and whole bodies of larvae.

To standardize data extraction, results from each study were categorized by organ type, as some studies analyzed multiple organs with varying miRNA responses (Bizuayehu et al., 2015; Y. Liu et al., 2024). The primary techniques employed across all studies were Small RNA sequencing (RNA-seq) for miRNA discovery and RT-qPCR for validation, ensuring consistency in miRNA quantification.

Across the 13 studies, there was considerable variability in the experimental conditions, particularly regarding temperature treatment and duration of heat stress exposure. Details of control and experimental groups varied by species, with range of temperature difference from 3°C to 13°C and exposure durations between 0 hours to 56 days. To address this heterogeneity, we grouped studies by species and thermal exposure conditions during the analysis phase.

Notably, four of the 13 studies originated from the same laboratory (J. Huang et al., 2018; Z. Liu et al., 2022; Ma et al., 2019; Zhao et al., 2023), raising potential concerns about data overlap. However, these studies utilized distinct fish species or experimental designs, justifying their inclusion. Furthermore, two papers authored by Sun et l., (2019a; 2019b) exhibited highly similar results, leading us to combine their data to avoid redundancy (Table 1).

**Table 1.** Summary of Experimental Designs from Studies Investigating the Role of microRNAs in Heat Stress Tolerance Across Various Fish Species. This table outlines the experimental setups from selected studies, specifying species, control group (CG) and treatment group (TG) temperatures, experimental conditions, and methods utilized (e.g., Small RNA sequencing and RT-qPCR). The table highlights the variability in temperature treatments and species used, facilitating a comparison of the methodologies applied to study microRNA expression in response to thermal stress in fish.

| Author, year | Species | CG temp/<br>°C | TG temp<br>/°C | Experimental design | Methods |
| --- | --- | --- | --- | --- | --- |
| Bizuayehu et al., 2015 | *Gadus morhua*<br>(Atlantic cod) | 4/9.5 | 4/9.5 | Embryos incubated in 8 incubators at 4°C and 8 at 9.5°C. 5 d post-hatching, 4°C juveniles in 4 incubators heated to 9.5°C, and 9.5°C juveniles in 4 incubators cooled to 4°C, at 0.9°C/d, 4 incubators were cooled to 4°C, respectively. The rising and cooling rates were 0.9°C/ d | Small RNA seq. and RT-qPCR |
| Qiang et al., 2017 | *Oreochromis niloticus*<br>(tilapia) | 28 | 35.5/36/3<br>6.5/37/37.<br>5/38/38.5<br>/39 | Acclimated at 28°C for 14 d, CG was kept at other temps for 4 d | Small RNA seq. and RT-qPCR |
| Zhang et al., 2017 | *Gymnocypris przewalskii*<br>(Tibetan naked carp) | 16 | 24 | The CG was maintained at 16°C. The TG was heated at 0.8°C/ h for 12 h at 24°C | Small RNA seq. and RT-qPCR |
| Bao et al., 2018 | *Oreochromis niloticus*<br>(tilapia) | 28 | 35 | Domestication at 28°C for three weeks was maintained at 28°C in the CG and 35°C in the TG. Samples were taken after 0, 6, 12, 24 and 48 h | Small RNA seq. and RT-qPCR |
| Huang et al., 2018 | *Oncorhynchus mykiss*<br>(rainbow trout) | 18 | 24 | Acclimated at 18°C for 7 d, the CG was kept at 18°C for 7 d, and the temps change rate in the TG was 1°C/d. Samples were taken after 7 d | Small RNA seq. and RT-PCR |
| Ma et al., 2019 | *Oncorhynchus mykiss*<br>(rainbow trout) | 18 | 24 | Acclimated at 18°C for 7 d, the CG was kept at 18°C for 7 d, and the temps change rate in the TG was 1°C/d. Samples were taken after 7 d | Small RNA seq. and RT-qPCR |
| Sun et al., 2019a | *Cyprinus carpio* L.<br>(normal carp) | 17 | 5、30 | The temp in the CG remained unchanged, and the temps change rate in the TG was 1°C/h, with a maximum change of 7°C/d. Samples were taken after 18 d | Small RNA seq. and qPCR |
| Sun et al., 2019b | *Cyprinus carpio* L.<br>(normal carp) | 17 | 5、30 | After domestication for 7 d, the CG was kept at 17°C, and the temps change rate in the TG was 1° C/h to the treatment temp, with a maximum of 7°C/d. Samples were taken after 18 d | Small RNA seq. and RT-qPCR |
| Vasadia et al., 2019 | *Trematomus bernacchii*<br>(Emerald rockcod) | -1.5 | 3.5 | Experimental temp. -1.5°C and 3.5°C were directly treated for 56 d, Samples were taken after 7/28/56 d | Small RNA seq. |
| Huang et al., 2022 | *Oncorhynchus mykiss*<br>(rainbow trout) | 16 | 26 | After 14 d of acclimation at 16°C, the CG continued to maintain 16°C, and the TG increased to 26°C at 1°C/d | Small RNA seq. and RT-qPCR |
| Liu et al., 2022 | *Oncorhynchus mykiss*<br>(rainbow trout) | 18 | 24 | CG was kept at 18°C for 7 d, and the temp. change rate in the TG was 1°C/d. Samples were taken after 7 d | Small RNA seq. and RT-qPCR |
| Zhao et al., 2023 | *Oncorhynchus mykiss*<br>(rainbow trout) | 18 | 24 | In vitro hepatocyte were cultured in a flask at test temp for 7 d | Small RNA seq. and RT-qPCR |
| Liu et al., 2024 | *Alosa sapidissima*<br>(American shad ) | 27 | 24、30 | Test temps directly treated for 3 d | Small RNA seq. and RT-qPCR |

### MicroRNAs and Enrichment Analysis

To further investigate the biological relevance of the identified miRNAs, a pathway enrichment analysis was performed on miRNAs that were recurrent across at least two studies. To streamline comparisons, species-specific indicators and suffixes subsequent to the contrasting arm were omitted. These miRNAs were converted to human miRNA orthologs to facilitate subsequent analysis (see Supplementary Table 3). The converted miRNAs were then analyzed using DIANA mirPath v.3 (Vlachos et al., 2015), a tool designed to elucidate the perturbed biological pathways influenced by these miRNAs. Within DIANA mirPath v.3, the target genes of these miRNAs were predicted using the microT-CDS v5.0 algorithm, which integrates predictive algorithms and meta-analytical fusion. The subsequent pathway analysis utilized Kyoto Encyclopedia of Genes and Genomes (KEGG) and Gene Ontology (GO) databases, with targets derived from TarBase v7.0. This dual approach allowed for the elucidation of pathway interactions and gene-level functional annotations (Blödorn et al., 2021).

To ensure robust results, the enrichment analysis was rigorously assessed using a false discovery rate (FDR)-adjusted p-value, with statistical significance evaluated via Fisher’s Exact Test. For both KEGG and GO analyses, a microT score threshold of 0.9 was applied, with an FDR-adjusted p-value threshold of less than 0.05 to filter for significantly enriched pathways and gene functions (Fig. 2). We used Cytoscape to create a targeted gene map (Shannon et al., 2003)(Fig. 3). These results provided insights into the primary cellular processes affected by heat stress in fish, which included metabolic and stress response pathways.

**Fig. 2.**
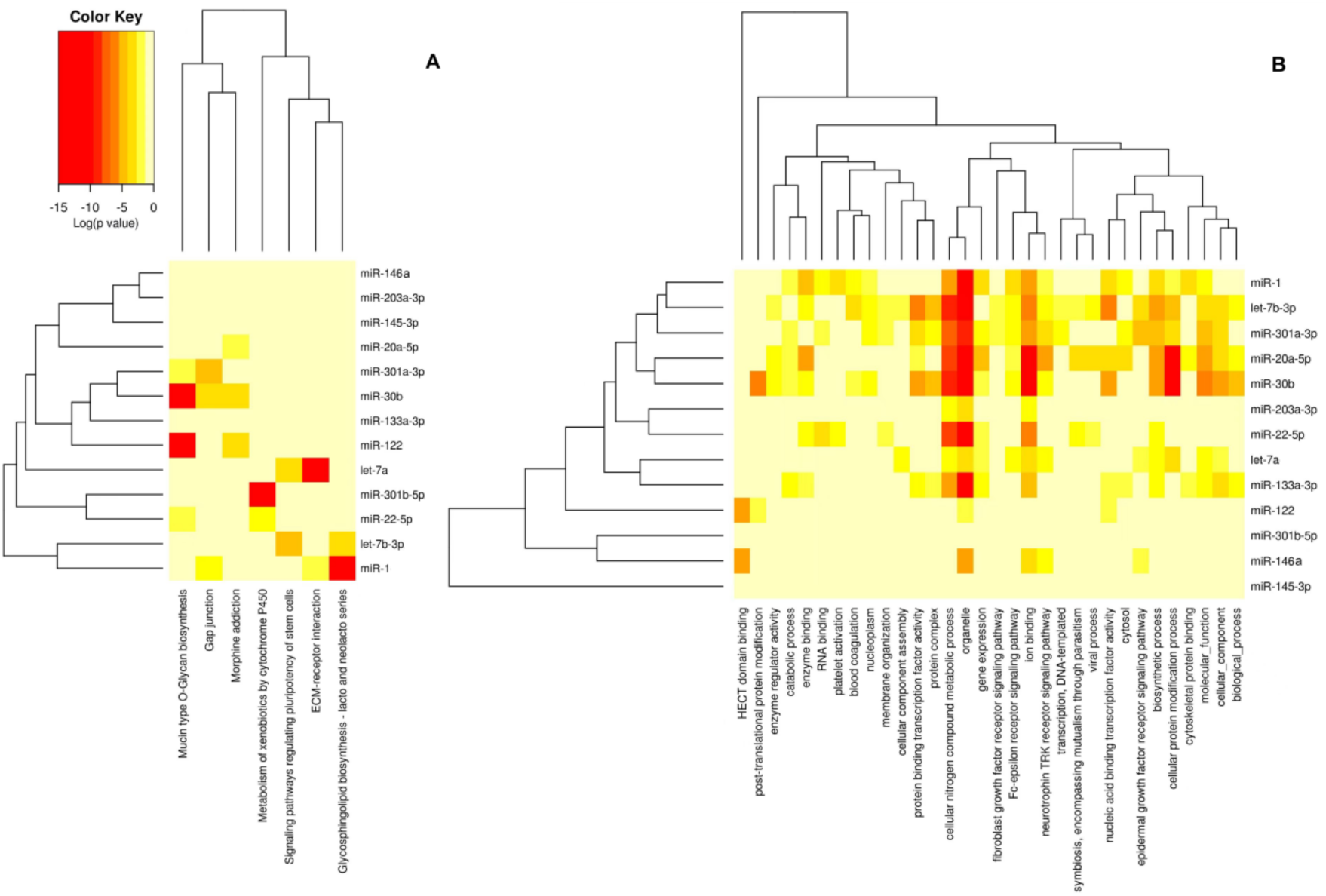
KEGG pathway (A) and GO term (B) enrichment analysis of recurring microRNAs identified in heat stress studies. The heatmaps display the most significant microRNAs involved in key biological pathways (KEGG) and gene ontology terms (GO), with the color scale indicating the fold enrichment. Red areas represent highly enriched microRNAs, indicating their significant role in heat stress responses across different species and experimental conditions. Hierarchical clustering further highlights the relationships between microRNAs and their involvement in these pathways and biological processes.

**Fig. 3.**
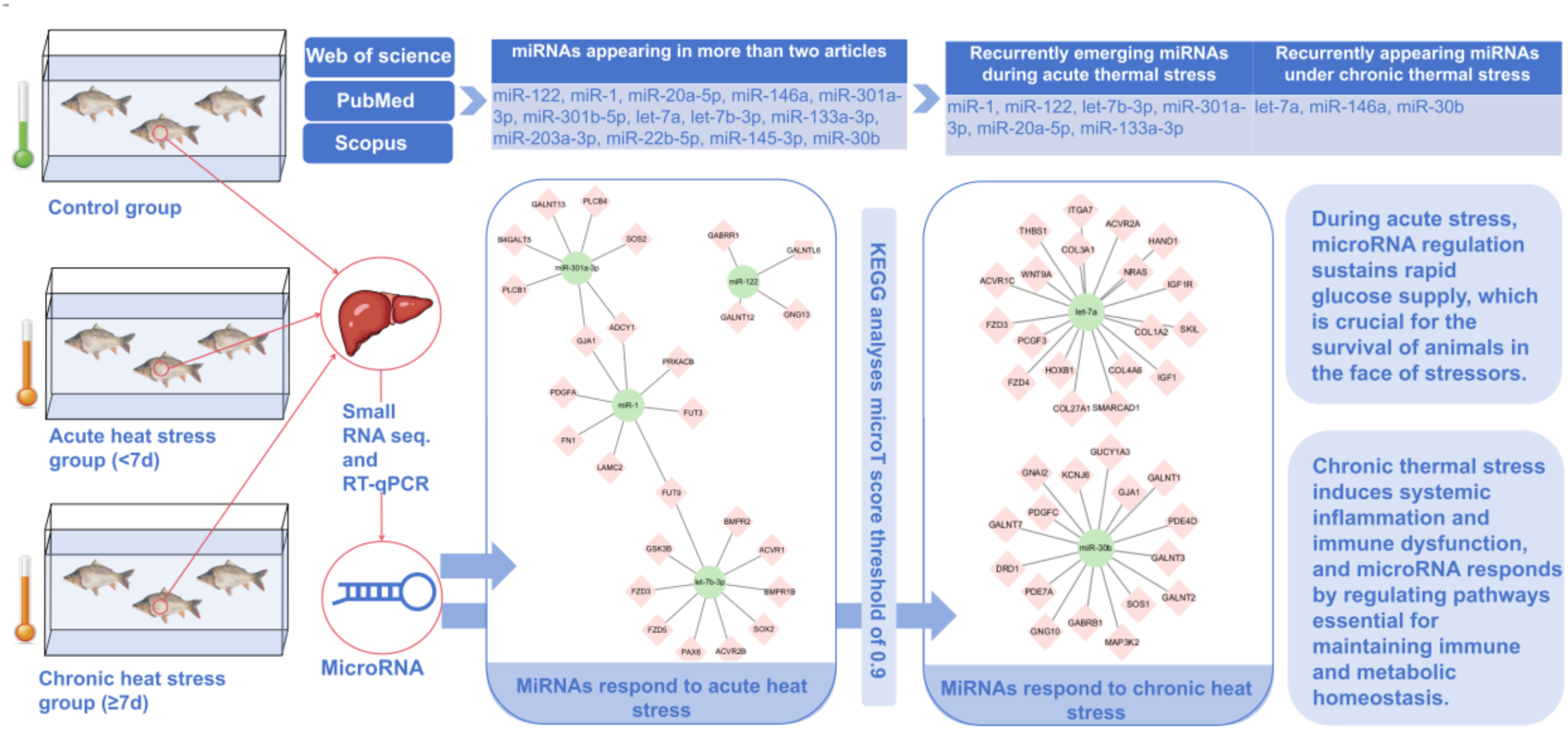
Heat responsiveness of microRNAs in fish under thermal stress: adaptive mechanisms and metabolic regulation. The duration of heat stress incorporated into the article is categorized into < 7 days (defined as acute heat stress) and ≥ 7 days (defined as chronic heat stress). Among the recurrent miRNAs in acute heat stress, the re-identified miRNAs are *miR-1*, *miR-122*, *let-7b-3p, miR-301a-3p*, *miR-20a-5p*, and *miR-133a-3p*, while for chronic heat stress, they are *let-7a*, *miR-146a*, and *miR-30b*. A target gene map was generated using Cytoscape for the target genes predicted by KEGG enrichment analysis with a microT score threshold of 0.9. The miRNAs responding to acute heat stress primarily regulate glucose homeostasis in fish, whereas those responding to chronic heat stress maintain immune and metabolic homeostasis.

## Results

### Differential microRNA of heat stress

As depicted in Table 2, the compiled 13 literature sources reveal a total of at least 214 significantly up-/down-regulated differential genes, comprising 122 upregulated and 113 downregulated genes. Notably, 15 of these genes are recurrent in at least two studies, with 12 appearing in both up- and 13 in down-regulation lists, among which 10 overlap in both categories and are highlighted in bold in Table 2. *miR-1* and *miR-1-3p* sequences are identical, so they are combined. Specifically, under heat stress, 11 miRNAs are upregulated: *miR-122, miR-1, miR-20a-5p, miR-146a, miR-301b-5p, let-7a, let-7b-3p, miR-7132b-5p, miR-133a-3p, miR-203a-3p, miR-22b-5p*. Conversely, 12 miRNAs are downregulated: *miR-122, miR-1, miR-20a-5p, miR-146a, miR-301a-3p, let-7a, let-7b-3p, miR-133a-3p, miR-203a-3p, miR-22b-5p, miR-145-3p, miR-30b*. In this research, heat stress lasting seven days or less is designated as acute stress, whereas heat stress exceeding seven days is classified as chronic stress. Among the 13 repeated differentially expressed miRNAs, *miR-1*, *miR-122*, *miR-7132b-5p*, *let-7b-3p*, *miR-301a-3p*, *miR-20a-5p*, and *miR-133a-3p* are miRNAs that repeat in acute heat stress, while *let-7a*, *miR-146a*, and *miR-30b* are miRNAs that repeat in chronic heat stress (Supplementary Table 4).

**Table 2.**
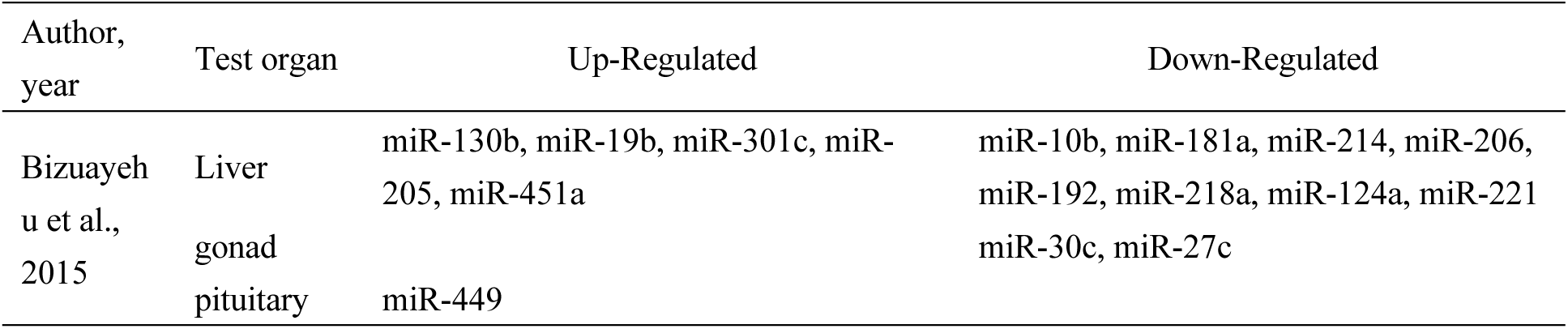
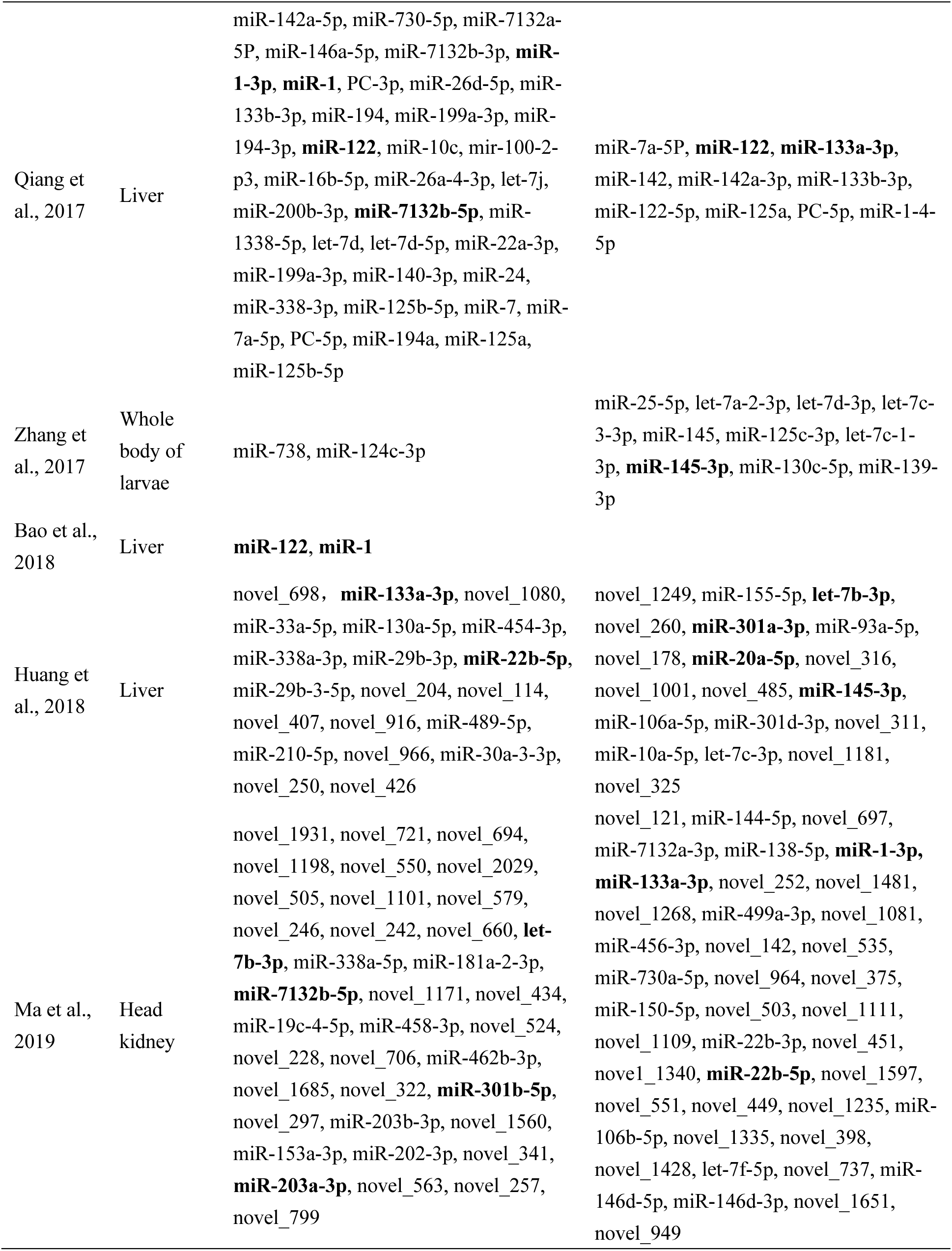
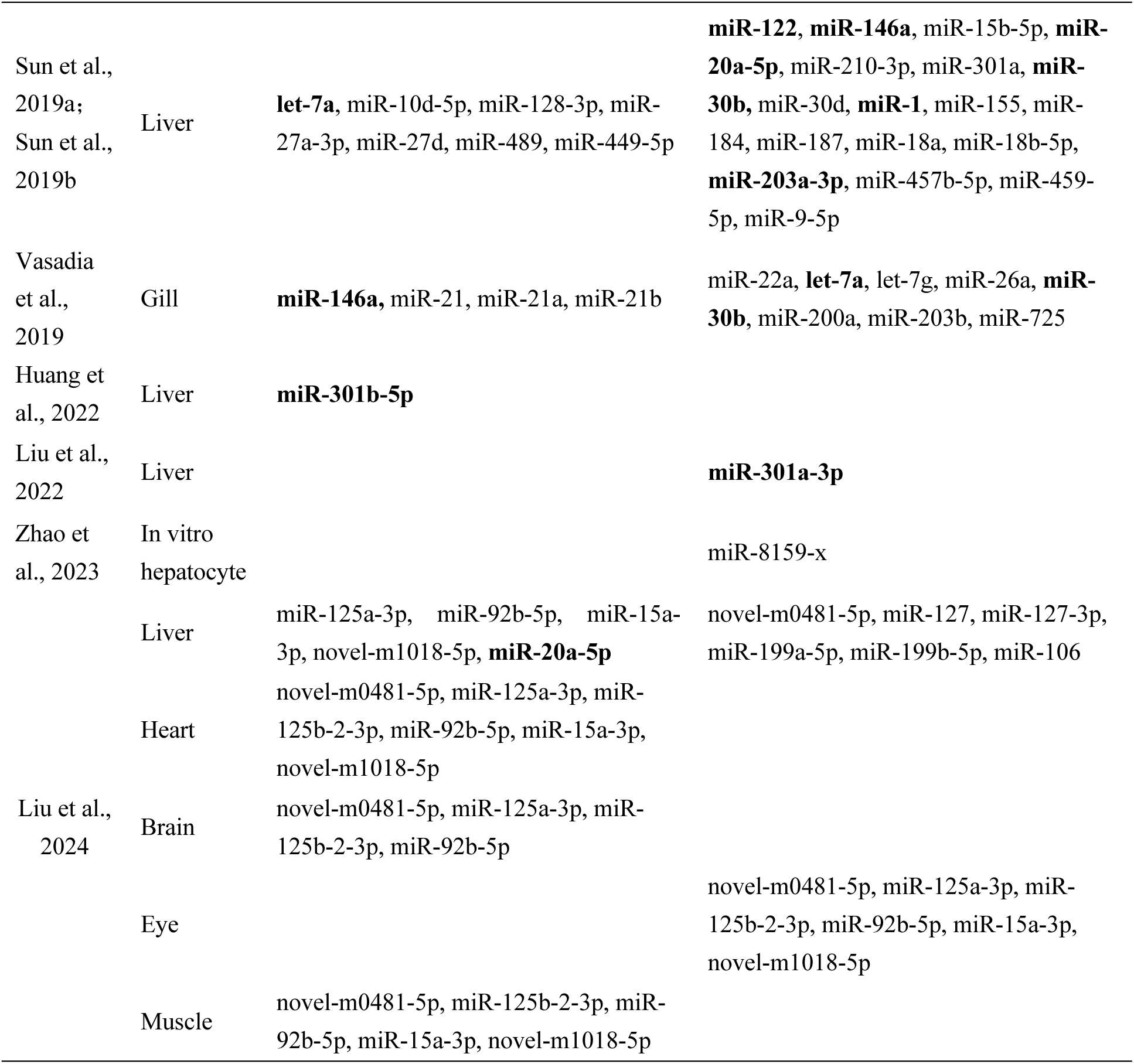
Differential expression of up-regulated and down-regulated miRNAs under heat stress in different tissues. The table outlines the differential up-regulated and down-regulated miRNAs in the organs examined in the selected studies, where the miRNAs in bold font are the miRNAs recurring in the two studies.

### MicroRNAs and enrichment analysis

KEGG pathway enrichment and GO functional analysis were performed on the selected 14 miRNAs that repeatedly emerged. However, due to DIANA mirPath v.3’s inability to recognize *ssa-miR-7132b-5p*, this miRNA was excluded from the KEGG and GO analyses, leaving 13 miRNAs for analysis (Fig. 2, Supplementary table 5). The KEGG and GO analyses revealed that the top 7 enriched pathways were: 1) *miR-122* targets GALNTL6, GALNT12, *miR-22-5p* targets GALNTL6, *miR-30b* targets GALNT7, GALNT1, GALNT3, GALNT2, *miR-301a-3p* targets B4GALT5, and GALNT13 regulates the Mucin type O-Glycon biosynthesis pathway. 2) *miR-1* targets ADCY1, GJA1, PRKACB, PDGFA, *miR-30b* targets GUCY1A3, DRD1, SOS1, PDGFC, GNAI2, GJA1, MAP3K2, *miR-301a-3p* targets ADCY1, SOS2, PLCB1, GJA1, PLCB4 regulates the Gap junction pathway. 3) *miR-122* targets GNG13, GABRR1, *miR-20a-5p* targets DRD1, PDE1B, GABBR2, KCNJ6, GNB5, *miR-30b* targets DRD1, PDE4D, GNG10, KCNJ6, GABRB1, GNAI2, PDE7A regulates Morphine addiction pathways. 4) *miR-22-5p* targets GSTM2, *miR-301b-5p* targets GSTO2 to regulate Metabolism of xenobiotics by cytochrome P450 pathway. 5) *let-7b-3p* targets GSK3B, FZD5, PAX6, BMPR1B, FZD3, ACVR1, ACVR2B, SOX2, BMPR2, *miR-30b* targets GUCY1A3, DRD1, SOS1, PDGFC, GNAI2, GJA1, MAP3K2, *let-7a* targets NRAS, HOXB1, HAND1, SMARCAD1, IGF1R, FZD3, FZD4, SKIL, ACVR2A, ACVR1C, IGF1, PCGF3, WNT9A regulates Signaling pathways regulating pluripotency of stem cells. 6) *miR-1* targets FN1, LAMC2, and *let-7a* targets THBS1, COL27A1, COL3A1, COL1A2, ITGA7, COL4A6 to regulate the ECM-receptor interaction pathway. 7) *miR-1* and *let-7b-3p* target FUT3, while FUT9 regulates the glycophingolipid biosynthesis lacto and neolacto series pathway (Fig. 2, Fig. 3). GO enrichment analysis reveals that the biological processes and metabolism during thermal stress in fish mainly encompass cellular, intracellular, and gene-level aspects, with cellular and transcriptome response functions to stimuli playing pivotal roles.

The KEGG enrichment analysis revealed that *miR-122*, *miR-22-5p*, *miR-30b*, and *miR-301a-3p* are significantly associated with the Mucin type O-Glycan biosynthesis pathway; *miR-1*, *miR-30b*, and *miR-301a-3p* are correlated with the Gap junction pathway; *miR-122*, *miR-20a-5p*, and *miR-30b* are significantly related to the Morphine addiction pathway; *miR-22-5p* and *miR-301b-5p* are associated with the Metabolism of xenobiotics by cytochrome P450 pathway; *let-7b-3p* and let-7a are significantly linked to the Signaling pathways regulating pluripotency of stem cells pathway; *miR-1* and *let-7a-5p* are significantly associated with the ECM-receptor interaction pathway; *miR-1* and *let-7b-3p* are significantly related to the Glycosphingolipid biosynthesis - lacto and neolacto series pathway.

The GO enrichment analysis indicates that the 13 recurring miRNAs are associated with various anabolic processes and functions, with the highest enrichment scores observed in cellular nitrogen compound metabolic processes, cellular organelles, ion homeostasis, neurotrophin TRK receptor signaling pathway, biosynthetic processes, and cellular protein modification processes. Among them, *miR-1*, *let-7b-3p*, *miR-301a-3p*, *miR-20a-5p*, *miR-30b*, *let-7a*, and *miR-133a-3p* are the most influential miRNAs.

## Discussion

The heat responsiveness of miRNAs in fish provides critical insights into their adaptive strategies amidst the growing threat of global warming. In our analysis, DE (Differentially expressed) miRNAs accounted for 7% of the total miRNAs investigated, reflecting variations in replication across the studies. KEGG pathway analysis revealed significant enrichment of DE miRNAs in systems related to metabolism, immune regulation, and biological processes, highlighting their multifaceted roles in thermal stress adaptation.

### MicroRNAs in glucose homeostasis and energy provisioning

Our analysis show that several miRNAs play crucial roles in regulating glucose homeostasis and energy provisioning in fish under heat stress. Notably, *let-7b-3p* and *let-7a* target multiple genes, exerting negative regulation over diverse biosynthetic and catabolic pathways by modulating glycogen synthase phosphorylation and inactivation. This influences hormone-regulated glucose homeostasis, Wnt signaling, transcription factors, and microtubule regulation. In bone marrow-derived osteoblasts (PyMS), these miRNAs enhance glucose transport, even at much lower concentrations significantly lower than those required by insulin (Mouchiroud, Eichner, Shaw, & Auwerx, 2014). Through this regulatory activity, *let-7b-3p* and *let-7a* efficiently rebalance cellular energy demands, reprogramming cellular functions in response to meet the increasing ATP needs imposed by thermal stress (Garcia & Shaw, 2017).

Previous studies have highlighted that the *let-7* family targets PRKAa1 (AMPKa1), a catalytic AMPK subunit of AMP-activated protein kinase (AMPK), which is essential for energy homeostasis, particularly in spinal cord function, where it is highly conserved systems such as the spinal cord (Craig, Trudeau, & Moon, 2014). When intracellular ATP levels decrease, AMPK inhibits ATP-consuming biosynthetic processes while activating ATP-regenerating pathways through phosphatase-mediated macromolecule breakdown. Under heat stress, fish experience metabolic disruptions, with glycogen/glucose mobilization becoming critical to meet elevated energy demands (Bacca et al., 2005; Oliveira, Rossi, Kucharski, & Da Silva, 2004). The coordinated actions of *let-7b-3p* and *let-7a* on these pathways allow rapid adaptation to such metabolic imbalances.

The miRNA, *miR-30b* further underscores the complexity of metabolic regulation, showing a positive correlation with phosphofructokinase liver isoform A (PFKLA), a key enzyme in cellular metabolism. The downregulation of *miR-30b* has been linked to increased expression of PCK2, an essential mitochondrial counterpart enzyme essential for glucose homeostasis (Stark et al., 2014; Stark & Kibbey, 2014). Additionally, *miR-20a-5p* has been found to regulate glucose metabolism in the liver by targeting and inhibiting the expression of homeobox B13 (HOXB13), a gene involved in cancer cell proliferation, invasion, and migration (H. Liu, Wang, Shen, Wu, & Li, 2022; Y. Liu et al., 2024; Wu et al., 2018).

Among the most frequently cited miRNAs in 13 thermal stress studies, *miR-122* and miR-1 stand out, appearing in 13 studies. These miRNAs, along with *miR-30b* and *miR-20a-5p*, are tightly linked to metabolic pathways such as pyruvate metabolism (ko00620), glycolysis/gluconeogenesis (ko00010), and the citrate cycle (TCA cycle) (ko00020) (Bao et al., 2018; Sun, Liu, et al., 2019). In particular, *miR-1* targets enzymes FUT3 and FUT9, which are involved in glycolipid and oligosaccharide biosynthesis, contributing to antigen synthesis and cellular proliferation. These miRNAs also play roles in glycosphingolipid biosynthesis, particularly the lacto and neolacto series pathways. Lactate, an indicator of glycolytic flux, is a precursor for glucose and glycogen resynthesis via the Cori cycle (Y. J. Chen et al., 2016), a metabolic cycle that efficiently converts lactate back to pyruvate, which is then transformed into glucose in the liver (Rabinowitz & Enerbäck, 2020). Elevated hepatic lactate levels, which are indicative of increased anaerobic glycolysis, suggest a higher rate of energy liberation under thermal stress (Sun, Liu, et al., 2019).

During heat stress, miRNAs such as *let-7b-3p*, *let-7a*, *miR-1*, *miR-122*, *miR-20a-5p*, and *miR-30b* exhibit significant expression changes, modulating glucose metabolism and their homeostasis within fish by targeting key genes intimately involved in glucose metabolism, including crucial enzyme genes in the glycolytic pathway. This regulatory network ensures that fish can continuously access sufficient energy supplies under adverse environmental conditions, highlighting their molecular adaptability to thermal stress.

### MicroRNAs in immunomodulation

A spectrum of miRNAs also controls immunomodulation under heat stress. *miR-122*, a conserved liver-specific miRNA in vertebrates, is important in maintaining liver homeostasis by regulating cholesterol, glucose, iron, and lipid metabolism (Thakral & Ghoshal, 2015). In GIFT (Genetically Improved Farmed Tilapia), *miR-122* expression under heat stress alterations correlates with immune system status (Qiang, Tao, et al., 2017). Increased *miR-122* expression is often a sign of liver injury (Thakral & Ghoshal, 2015; K. Wang et al., 2009).

KEGG analysis revealed that *miR-122*, along with *miR-30b*, *miR-22-5p*, and *miR-301a-3p*, are central regulators of mucin-type O-glycan biosynthesis, essential for forming protective barriers in epithelial tissues (Brockhausen & Argüeso, 2020; Y. Zhang et al., 2021). Meanwhile, the mitogen-activated protein kinase (MAPK) cascade, a highly conserved pathway involved in numerous physiological processes, plays a vital role in immune responses (Mathien, Tesnière, & Meloche, 2021). Recent research has emphasized the connection between MAPK signaling and T-cell immunity in fish, under prolonged heat stress (Wei et al., 2020).

In response to thermal stress, fish adjust their immune defenses by regulating miRNAs such as *miR-122*, *miR-30b*, *miR-22-5p*, and *miR-301a-3p*, which in turn modulate pathways like mucin-type O-glycan biosynthesis. This coordinated molecular response facilitates tissue repair, strengthens immune barriers, and enhances overall resilience to heat-induced damage.

### MicroRNAs in heat shock proteins and physiological processes

miRNAs also play significant roles in modulating heat shock proteins (HSPs) and associated physiological processes. Previous studies have shown that miRNAs like *let-7b-3p*, *miR-301a-3p*, and *miR-20a-5p* target members of the HSP family (HSP90B2, HSP90BA, HSP90BB, HSP40), key molecules involved in the thermal stress response (J. Huang et al., 2018; Z. Liu et al., 2022; Ma et al., 2019). While this study did not detect a significant impact on HSPs, miRNAs such as *miR-122* and *miR-30b* were found to target genes related to critical physiological processes, including dopamine receptor signaling and second messenger systems like cAMP.

The miRNAs *let-7b-3p* and *let-7a* are also implicated in signaling pathways that regulate stem cell pluripotency, with aberrant expression potentially disrupting cellular functions (Bernstein, R, Nfonsam, & Bernstei, 2013). Furthermore, *let-7a*’s strong correlation with extracellular matrix (ECM) receptor interactions underscores its importance in cellular adaptation to environmental stress.

### MicroRNAs in acute and chronic heat stress

In studies differentiating between acute and chronic heat stress, several miRNAs are identified as being exceptionally responsive. Acute stress is marked by a rapid onset of severe metabolic disruptions, including insulin resistance, mediated by miRNAs such as *miR-1*, *miR-122*, *let-7b-3p*, *miR-301a-3p*, and *miR-133a-3p* . In contrast, chronic heat stress, which involves prolonged exposure, elicits systemic inflammation and immune dysregulation, with miRNAs like *let-7a*, *miR-146a*, and *miR-30b* playing pivotal roles (Fig. 3, Supplementary Table 5).

In response to acute stress, fish utilize responsive miRNAs to regulate glucose levels by targeting specific miRNAs, which in turn elevate blood glucose and stimulate ATP production, vital for energizing the immune system and internal organs. These miRNAs are crucial for the animal’s survival in the face of immediate stressors (Barreto & Volpato, 2006; Chan, Poller, Swirski, & Russo, 2023; L. Liu et al., 2024; Sharma, Akre, Chakole, & Wanjari, 2022).

Fish respond to chronic heat stress by modulating miRNAs that promote angiogenesis, regulate reactive oxygen species (ROS) signaling, and activate pathways essential for maintaining immune and metabolic homeostasis. These miRNAs contribute to wound healing, stress recovery, and overall survival during extended periods of thermal challenge (Cantet, Yu, & Ríus, 2021; Chan et al., 2023; Marik & Bellomo, 2013; Sejian, Bhatta, Gaughan, Dunshea, & Lacetera, 2018).

This study discusses recurring elements among numerous DE miRNAs, important in fish heat stress. As highly conserved miRNAs, they could serve as hepatic biomarkers for fish heat stress amidst global warming, monitoring wild fish populations’ health, enhancing selective breeding programs in aquaculture, or informing conservation strategies for vulnerable species. However, limitations persist. Firstly, disparate methods to induce heat stress in fish analyzed herein hinder a clear depiction of stress types and their relevance to global climate change. Secondly, the strong tissue specificity of miRNAs necessitates caution regarding sampling locations across studies. Given the liver’s preeminent vulnerability under stress, our discussion is confined to its response. Thermal stress responses in other tissues remain to be explored.

## Conclusions

Our review highlights the critical role of miRNAs in modulating physiological and immune responses to heat stress in fish, emphasizing their significance in adaptive mechanisms under changing environmental conditions. Through differential expression, miRNAs such as *let-7b-3p*, *let-7a*, *miR-1*, *miR-122*, and *miR-30b* were shown to regulate key metabolic pathways, including glucose homeostasis, glycolysis, and the TCA cycle, ensuring energy provision during thermal stress. Additionally, miRNAs involved in immunomodulation, such as *miR-122* and *miR-30b*, were linked to mucin-type O-glycan biosynthesis and MAPK signaling, further enhancing the defense mechanisms of the fish.

Although heat shock proteins were not significantly impacted in this study, miRNAs targeting HSP-related genes were implicated in broader physiological adaptations, including dopamine receptor signaling and ECM interactions. The miRNA profiles also revealed distinct responses to acute and chronic heat stress, with acute stress triggering insulin resistance and metabolic disruptions, while chronic stress led to immune dysregulation and systemic inflammation.

Hence, these findings provide a comprehensive understanding of how miRNAs mediate complex molecular responses to heat stress, offering insights into potential targets for enhancing thermotolerance in fish species as climate change intensifies.

## ACKNOWLEDGEMENTS

We thank Guangxi University for providing access to the literature search software.

## AUTHOR CONTRIBUTIONS

Conceptualization – LT, MM; Data curation – LT; Formal analysis – LT; Funding acquisition – MM; Investigation – LT, MM; Methodology - – LT, MM; Project administration – LT, MM; Resources – LT, MM; Software – N/A; Supervision – MM; Validation – LT; Visualization - – LT; Writing – original draft - – LT, MM; Writing – review and editing – LT, MM.

## DATA AVAILABILITY STATEMENT

The data that was compiled and analyzed are available at: DOI 10.5281/zenodo.13825048

## ETHICS STATEMENT

Since this study is a systematic literature review with analysis, hence there are no ethical concerns to report.

