## Supplementary Files for "Fish MicroRNA Responses to Thermal Stress: Insights and Implications for Aquaculture and Conservation Amid Global Warming"

**_____________________________________________________________________**

**Supplementary Table 1 Search language logic in three databases.** The table shows that the literature in this paper is derived from three databases in a Web of Science, PubMed, and Scopus using #1 microRNA OR miRNA, #2 heat OR high temperature OR stress, #3 tolerance OR resistance OR adaption, #4 fish These 4 groups of keywords are obtained by the combined search.

| Web of Science、PubMed and Scopus | #1 microRNA OR miRNA |
| --- | --- |
|  | #2 heat OR high temperature OR stress |
|  | #3 tolerance OR resistance OR adaption |
|  | #4 fish |

Search combination: #1 AND #2 AND #3 AND #4.

**Supplementary Table 2 Details of the topic in selected studies.** The table Outlines the author and year of the selected study and the corresponding paper title, which shows the focus of the research and the fish species tested. The reader can get some basic information about the selected document based on the information.

| No. | Author, year | Article title |
| --- | --- | --- |
| 1 | Bizuayehu et al., 2015 | Temperature during early development has long-term effects on microRNA expression in Atlantic cod |
| 2 | Qiang et al., 2017 | The expression profiles of miRNA-mRNA of early response in genetically improved farmed tilapia (*Oreochromis niloticus*) liver by acute heat stress |
| 3 | Zhang et al., 2017 | Integrated mRNA and microRNA transcriptome analyses reveal regulation of thermal acclimation in *Gymnocypris przewalskii*: A case study in Tibetan Schizothoracine fish |
| 4 | Bao et al., 2018 | Responses of blood biochemistry, fatty acid composition and expression of microRNAs to heat stress in genetically improved farmed tilapia (*Oreochromis niloticus*) |
| 5 | Huang et al., 2018 | Identification and characterization of microRNAs in the liver of rainbow trout in response to heat stress by high-throughput sequencing |
| 6 | Ma et al., 2019 | High-throughput sequencing reveals microRNAs in response to heat stress in the head kidney of rainbow trout (*Oncorhynchus mykiss*) |
| 7 | Sun et al., 2019a | Potential regulation by miRNAs on glucose metabolism in liver of common carp (*Cyprinus carpio*) at different temperatures |
| 8 | Sun et al., 2019b | Analysis of miRNA-seq in the liver of common carp (*Cyprinus carpio* L.) in response to different environmental temperatures |
| 9 | Vasadia et al., 2019 | Characterization of thermally sensitive miRNAs reveals a central role of the FoxO signaling pathway in regulating the cellular stress response of an extreme stenotherm, *Trematomus bernacchii* |
| 10 | Huang et al., 2022 | miR-301b-5p and its target gene nfatc2ip regulate inflammatory responses in the liver of rainbow trout (*Oncorhynchus mykiss*) under high temperature stress |
| 11 | Liu et al., 2022 | Gene ssa-miR-301a-3p improves rainbow trout (*Oncorhynchus mykiss*) resistance to heat stress by targeting hsp90b2 |
| 12 | Zhao et al., 2023 | Potential role of miR-8159-x in heat stress response in rainbow trout (*Oncorhynchus mykiss*) |
| 13 | Liu et al., 2024 | Integrated transcriptome and microRNA analysis reveals molecular responses to high-temperature stress in the liver of American shad (*Alosa sapidissima*) |

**Supplementary Table 3 Details of the comparison between the above 13 recurring miRNAs and human homologous miRNAs in two papers in the literature were selected**. This table shows repeated miRNAs names, prefixes, mature sequences, and changes in the included studies, corresponding to human miRNAs names and their mature sequences. The sequence difference between human homologous miRNAs and the mature miRNAs included in the study was less than or equal to 4. This paper will conduct KEGG and GO enrichment analysis based on human homologous miRNAs. Since miR-1 and Mir-1-3P have the same base sequence, we combined them. Because ssa-miR-7132a-3p does not have a less distinct human homologous miRNAs, there are no corresponding references, so it is excluded from the discussion. The base differences between repeat microRNA sequences and similar human microRNA sequences are all underlined in the table.

| MicroRNA | Species prefix | Mature sequence | Human microRNA | Mature sequence |
| --- | --- | --- | --- | --- |
| miR-122 ↑↓ | dre/ccr | UGGAGUGUGACAAUGGUGUUUG | hsa-miR-122-5p | UGGAGUGUGACAAUGGUGUUUG |
| miR-1 ↑↓ | dre/ccr | UGGAAUGUAAAGAAGUAUGUAU | hsa-miR-1-3p | UGGAAUGUAAAGAAGUAUGUAU |
| miR-20a-5p ↑↓ | ssa/ccr/pma | UAAAGUGCUUAUAGUGCAGGUAG | Hsa-miR-20a-5p | UAAAGUGCUUAUAGUGCAGGUAG |
| miR-146a ↑↓ | tbe/ccr | UGAGAACUGAAUUCCAUAGGUUGU | hsa-miR-146b-5p | UGAGAACUGAAUUCCAUAGGCUG |
| miR-301b-5p ↑ | ssa | GCUUUGACGAUGUUGCACUACU | hsa-miR-301b-5p | GCUCUGACGAGGUUGCACUACU |
| miR-301a-3p ↓ | ssa | CAGUGCAAUAGUAUUGUCAUAGC | hsa-miR-301a-3p | CAGUGCAAUAGUAUUGUCAAAGC |
| let-7a ↑↓ | tbe/ccr | UGAGGUAGUAGGUUGUAUAGUU | hsa-let-7a-5p | UGAGGUAGUAGGUUGUAUAGUU |
| let-7b-3p ↑↓ | ssa | CUGUACAACCUACUGCCUUCCC | hsa-let-7b-3p | CUAUACAACCUACUGCCUUCCC |
| miR-133a-3p ↑↓ | dre/ssa | UUUGGUCCCCUUCAACCAGCUG | hsa-miR-133a-3p | UUUGGUCCCCUUCAACCAGCUG |
| miR-145-3p  ↓ | ssa | AUUCCUGGAAAUACUGUUCUU | hsa-miR-145-3p | GGAUUCCUGGAAAUACUGUUCU |
| miR-203a-3p ↑↓ | ssa/ccr | GUGAAAUGUUUAGGACCACUUG | hsa-miR-203a-3p | GUGAAAUGUUUAGGACCACUAG |
| miR-22b-5p ↑↓ | ssa | CGUUCUUCACUGGCUAGCUUU | hsa-mir-22-5p | AGUUCUUCAGUGGCAAGCUUUA |
| miR-30b ↓ | ccr/tbe | UGUAAACAUCCUACACUCAGCU | hsa-mir-30b-5p | UGUAAACAUCCUACACUCAGCU |

**Supplementary Table 4 Recurrent up-regulation and down-regulation of miRNAs in acute heat stress and chronic heat stress were studied in two studies.** Acute heat stress and chronic heat stress were divided according to whether the duration of heat stress in the study exceeded 7 days.

| **Stress type** | **Author, year** | **Up-Regulated** | **Down--Regulated** |
| --- | --- | --- | --- |
| Acute stress | Bizuayehu et al., 2015 | / | / |
|  | Qiang et al., 2017 | miR-1, miR-122, miR-7132b-5p | miR-122, miR-133a-3p |
|  | Zhang et al., 2017 |  | miR-145-3P |
|  | Bao et al., 2018 | miR-1, miR-122, |  |
|  | Huang et al., 2018 | miR-133a-3p, miR-22b-5p | let-7b-3p, miR-301a-3p, miR-20a-5p |
|  | Ma et al., 2019 | let-7b-3p, miR-7132b-5p | miR-145-3p, miR-133a-3p |
|  | Liu et al., 2022 |  | miR-301a-3p |
|  | Zhao et al., 2023 |  | miR-8159-x |
|  | Liu et al., 2024 | miR-20a-5p |  |
| Chronic stress | Sun et al., 2019a; 2019b | let-7a | miR-122, miR-146a, miR-1, miR-20a-5p, miR-30b, miR-203a-3p |
|  | Vasadia et al., 2019 | miR-146a | let-7a, miR-30b |
|  | Huang et al., 2022 | miR-301b-5p |  |

**Supplementary Table 5 Target genes and KEGG pathways predicted by** **mirPath v.3 of 13 recurring miRNAs in** **DIANA TOOLS.**

| **KEGG pathway** | **miRNAs** | **Genes** |
| --- | --- | --- |
| Mucin type O-Glycan biosynthesis | miR-122 | GALNTL6 |
|  |  | GALNT12 |
|  | miR-22-5p | GALNTL6 |
|  | miR-30b | GALNT7 |
|  |  | GALNT1 |
|  |  | GALNT3 |
|  |  | GALNT2 |
|  | miR-301a-3p | B4GALT5 |
|  |  | GALNT13 |
| Glycosphingolipid biosynthesis - lacto and neolacto series | miR-1 | FUT3 |
|  |  | FUT9 |
|  | let-7b-3p | FUT9 |
| Metabolism of xenobiotics by cytochrome P450 | miR-22-5p | GSTM2 |
|  | miR-301b-5p | GSTO2 |
| ECM-receptor interaction | miR-1 | FN1 |
|  |  | LAMC2 |
|  | let-7a-5p | THBS1 |
|  |  | COL27A1 |
|  |  | COL3A1 |
|  |  | COL1A2 |
|  |  | ITGA7 |
|  |  | COL4A6 |
| Gap junction | miR-1 | ADCY1 |
|  |  | GJA1 |
|  |  | PRKACB |
|  |  | PDGFA |
|  | miR-30b | GUCY1A3 |
|  |  | DRD1 |
|  |  | SOS1 |
|  |  | PDGFC |
|  |  | GNAI2 |
|  |  | GJA1 |
|  |  | MAP3K2 |
|  | miR-301a-3p | ADCY1 |
|  |  | SOS2 |
|  |  | PLCB1 |
|  |  | GJA1 |
|  |  | PLCB4 |
| Signaling pathways regulating pluripotency of stem cells | let-7b-3p | GSK3B |
|  |  | FZD5 |
|  |  | PAX6 |
|  |  | BMPR1B |
|  |  | FZD3 |
|  |  | ACVR1 |
|  |  | ACVR2B |
|  |  | SOX2 |
|  |  | BMPR2 |
|  | let-7a-5p | NRAS |
|  |  | HOXB1 |
|  |  | HAND1 |
|  |  | SMARCAD1 |
|  |  | IGF1R |
|  |  | FZD3 |
|  |  | FZD4 |
|  |  | SKIL |
|  |  | ACVR2A |
|  |  | ACVR1C |
|  |  | IGF1 |
|  |  | PCGF3 |
|  |  | WNT9A |
| Morphine addiction | miR-122 | GNG13 |
|  |  | GABRR1 |
|  | miR-20a-5p | DRD1 |
|  |  | PDE1B |
|  |  | GABBR2 |
|  |  | KCNJ6 |
|  |  | GNB5 |
|  | miR-30b | DRD1 |
|  |  | PDE4D |
|  |  | GNG10 |
|  |  | KCNJ6 |
|  |  | GABRB1 |
|  |  | GNAI2 |
|  |  | PDE7A |
